# Canopy1/Cnpy1 is required for proper V2R processing and transport; its loss impairs the function and circuit organization of basal/V2R vomeronasal sensory neurons

**DOI:** 10.1101/2025.11.19.689260

**Authors:** Nicholas Mathias, Nikki Dolphin, Anthony Ruquet, Paolo E Forni

## Abstract

The vomeronasal organ (VNO) is a specialized chemosensory structure in the nasal cavity that detects pheromones and mediates social and reproductive behaviors. The VNO of rodents is populated by different types of vomeronasal sensory neurons (VSNs). Apical VSNs, located near the lumen, express the transcription factor (TF) Meis2, V1R family receptors, and the G protein subunit Gao; the VSNs distributed closer to the basal lamina express the TF Tfap2e/AP-2ε, V2R receptors, and the G protein subunit Gai2. In addition, sparse cells in the VNO express the Formyl Peptide Receptors (FPRs). Single-cell mRNA sequencing (scRNA Seq) identified over 980 differentially expressed genes between these cell types, with many linked to the endoplasmic reticulum (ER). Among these ER proteins, Canopy1 (Cnpy1), was found to be among the most enriched genes in V2R+ VSNs. Previously studied only in zebrafish, Cnpy1 was found to affect Fgfr1 signaling and is thus also known as “FGF signaling regulator-1”. In a previous study, we discovered that AP-2e upregulates Cnpy1 expression. Although Cnpy1 knockout mice are viable and have normal VNO development at birth, they experience a progressive degeneration and loss of V2R-expressing VSNs. Prior to symptoms, the basal VSNs of KO mice display reduced V2R protein immunoreactivity in the soma and a complete absence of the protein at the lumen of the VNO, rendering the neurons non-functional. Cnpy1 KOs exhibit altered guidance cues and adhesion molecule expression, along with disrupted connectivity to the accessory olfactory bulb (AOB). Our study shows that distinct neuronal types depend on unique ER protein repertoires to maintain proper proteostasis. The loss of Cnpy1 highlights the importance of cell–type–specific ER factors in the differentiation and function of specific neurons, revealing mechanisms that drive neuronal diversity and vulnerability to ER gene disruption.

## Introduction

Different types and subtypes of cells are defined by their unique set of proteins expressed. Which proteins a cell can successfully express result from complex transcriptional, post-transcriptional, translational, and post-translational controls and modifications (Ellgaard et al., 2016; Sun and Brodsky, 2019). High-throughput single-cell transcriptomics is leading to an unprecedented understanding of cellular diversity. However, while transcriptomics enables us to distinguish and analyze cells based on gene expression, cellular differences in translational and post-translational processing remain mostly unexplored. While general proteostatic mechanisms are proposed, the diversity of ER composition across different cells is largely overlooked as a key variable in understanding development, aging, and neuronal degeneration.

The sensory epithelium of the rodent VNO is primarily populated by vomeronasal sensory neurons (VSNs), which either express V1R or V2R receptor gene families or Formyl Peptide Receptors (FPRs) (Ackels et al., 2014; Dietschi et al., 2013; Liberles et al., 2009; Riviere et al., 2009). The V1R and V2R VSNs both arise from a shared pool of progenitors, which in postnatal animals are primarily found in the marginal zone. The VNO, like the main olfactory epithelium, exhibits postnatal neurogenesis throughout life (Brann et al., 2014). After differentiation, V1R VSNs localize apically toward the lumen, while V2R VSNs localize toward the basal lamina. The V1R/V2R dichotomy is determined by differential Notch signaling in newly formed VSNs (Katreddi et al., 2022), after which these neuronal types begin to diverge in their expression of transcription factors, G protein subunits, adhesion molecules, specific VR genes, and guidance receptors (Hills et al., 2024; Katreddi et al., 2022; Lin et al., 2022). V1R- and V2R-expressing VSNs project and synapse onto projection neurons of the anterior and posterior AOB, respectively (Fig. 1A). The projection neurons in these two AOB regions arise from distinct embryonic lineages (Huilgol et al., 2013).

**Figure 1.**
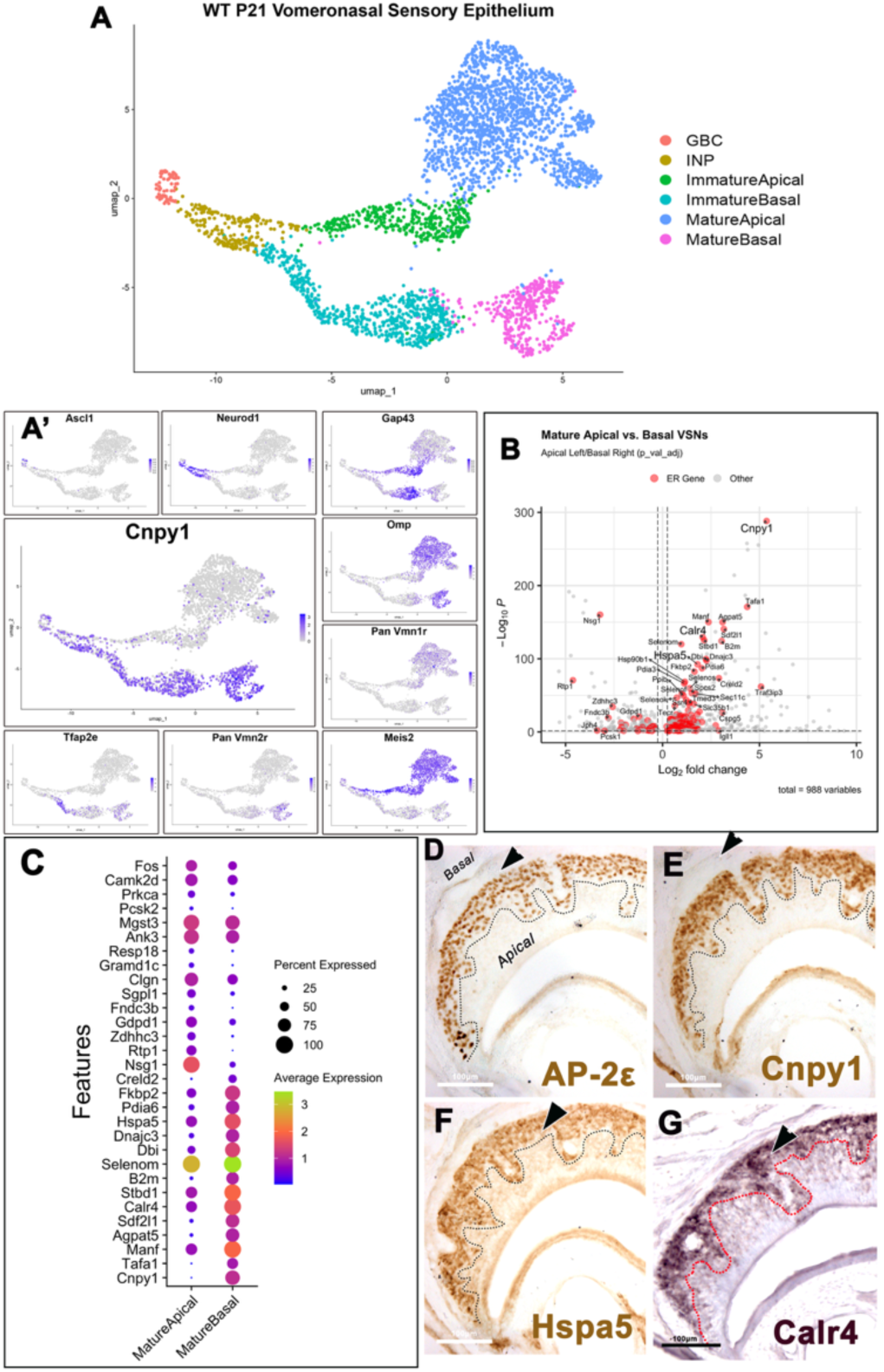
Transcriptomic and histological characterization of apical and basal VSNs. **(A)** UMAP of WT P21 VNO single-cell RNA-seq showing progression from *Ascl1*⁺ GBCs and *Neurod1*⁺ INPs to *Gap43*⁺ immature and mature apical (*V1R*) and basal (*V2R*) VSNs. (A′) Feature plots for key markers, including *Ascl1*, *Neurod1*, *Gap43*, *Omp*, *Tfap2e*, pan-*Vmn1r*, pan-*Vmn2r*, *Meis2*, and the ER gene *Cnpy1*, note its enrichment along the basal VSNs from immature to mature. (B) Volcano plot of mature apical vs. basal VSNs highlighting ER-enriched genes (*Cnpy1*, *Hspa5*, *Calr4*). (C) Dot-plot of the top ER-related differentially expressed genes based on GO analysis. (D–F) Immunohistochemistry for AP-2ε, Cnpy1, and HSPA5/BiP confirms basal enrichment. (G) *In situ* hybridization anti *Calr4* shows strong basal expression. Scale bars: 100μm.

Recent papers have uncovered multiple previously unknown genetic differences across the two main neuronal types of VSNs, suggesting that these cells are more diverse than previously known (Devakinandan and Dani, 2024; Hills et al., 2024; Katreddi and Forni, 2021; Katreddi et al., 2022; Lin et al., 2022; Naik et al., 2020).

We previously reported that the transcription factor AP-2ε, which is among the most differentially expressed genes between V1R+ and V2R+ VSNs, is essential to maintain the basal VSN program after it is initiated by Notch signaling and Bcl11b expression (Enomoto et al., 2011; Katreddi et al., 2022). CUTCRUN data from mice with ectopic AP-2ε expression in V1R neurons indicate that AP-2ε functions as a master regulator with both transcriptional activation and repression functions. In the VNO, AP-2ε upregulates the expression of about one-third of the basal (V2R) neuronal type-specific genes while repressing roughly one-third of the apical (V1R) neuronal type genes (Lin et al., 2022). Ectopic expression of AP-2ε in the V1R+/apical VSNs was found to enhance the expression of the basally enriched ER protein Cnpy1, the Golgi protein Selenom, along with the ER protein Calreticulin-4, which has been proposed to play a key role in V2R receptor post-translational processing. Repressor activity has been observed for the V1R/apical enriched ER protein Calreticulin (Dey and Matsunami, 2011; Lin et al., 2022)

Apical and basal VSNs show significant differences in ER structure and expression of ER protein repertoires, respectively (Devakinandan and Dani, 2024). These findings suggest that the two major VSN neuronal types rely on distinct, cell-type-specific ER protein molecular machinery to sustain their specialized proteostasis. Consequently, if each neuronal subtype relies on its own unique ER protein repertoire, this could lead to different genetic vulnerabilities across chemosensory neuronal populations when ER proteins are disrupted.

Cnpy1 is the founder of the canopy family (Li et al., 2025). The canopy proteins are part of the larger saposin-like superfamily, and all share the common saposin fold, which is believed to facilitate their interaction with plasma membranes and enable dimerization (Schildknegt et al., 2019). The only existing in vivo studies in zebrafish pointed to roles for Cnpy1 in establishing the mid-hindbrain barrier with a proposed role in Fgfr1/Fgf8 signaling (Hirate and Okamoto, 2006; Matsui et al., 2011). In this study, we examined Cnpy1 expression in the VNO of mice and the impact of Cnpy1 loss-of-function. Our work, along with an independent simultaneous study (Devakinandan et al., 2025), reveals a previously unknown role for Cnpy1 as an indispensable part of the gene regulatory networks for V2R VSNs and is a central component of their cell-type-specific ER protein repertoire.

## Results

### Endoplasmic Reticulum Proteins of the VNO Exemplify VSN Identity Differences and Maturation

Single-cell suspensions were harvested from WT P21 mice, sequenced, and processed using Seurat. The resulting cells were subset to isolate the sensory epithelium of the VNO to generate a UMAP capturing the major stages of VSN differentiation (Fig. 1A). This analysis resolved Ascl1⁺ globose basal progenitors (GBCs), their transition into Neurog1⁺/NeuroD1⁺ immediate neuronal precursors (iNPs), followed by more advanced neuronal precursors in which Neurod1 expression falls as GAP43 rises. These cells then differentiate into GAP43⁺ immature VSNs and ultimately mature into OMP⁺ VSNs along either the Meis2/V1R or AP-2e/V2R trajectories. Together, these data recapitulate the expected developmental continuum from progenitor to fully differentiated V1R- or V2R-expressing sensory neurons endowed with mature sensory and synaptic competence. We performed a Wilcox Rank-Sum differential gene expression analysis between mature basal (V2R) and apical (V1R) VSNs. In line with a recent study (Devakinandan and Dani, 2024), we observed a significant representation of genes coding for proteins belonging to the endoplasmic reticulum (ER) or which control ER biosynthesis and functions according to the “Endoplasmic Reticulum” Gene Ontology (GO) term (GO:0005783). In mature apical (V1R/Meis2+) VSNs, 31 of the 279 genes that are enriched in apical VSNs (11.1%) were classified as ER-related genes. In mature basal VSNs (V2R/AP-2e+), 145 of the 709 genes enriched in basal VSNs (20.5%) were identified as ER-related proteins. To visualize such diversity in ER protein expression across mature cell types, we generated a dot plot (Fig. 1C) showing the top 15 ER-related genes for apical and basal VSNs. Among the various enriched genes in the V2R VSNs, we noticed strong expression of Hspa5, Canopy1 (Cnpy1), and Calreticulin4, (Fig. 1A-C). Immunolabeling and in situ hybridization confirmed enrichments of ER genes in the basal VSNs (Fig.1 D-G).

#### Cnpy1 is the only member of the Cnpy family enriched in V2R VSNs

Feature plot analysis of Cnpy1 mRNA expression revealed that the Cnpy1 gene starts to be expressed as early as in the proliferative GBCs (Ascl1+) (Fig. 1A; Fig. 2B) and in early progenitor precursors (Neurog1-Neurod1). However, its mRNA expression rapidly declines in differentiating/maturing (Meis2+; Gai2/Gnai2) V1R VSNs while remaining active and highly correlated with the expression of genes belonging to the V2R basal lineage (Fig. 1A, 2A). Correlation plot confirmed the positive correlation with maturation of basal/V2R+ VSN as they express Tfap2e (AP-2e), Gao (Gnao1) and V2Rs (Fig.2A). Notably, other members of the Cnpy family (Schildknegt et al., 2019), such as Cnpy2, Cnpy3, and Cnpy4, and Cnpy5/Mzb1 were also found to be variably expressed. While Cnpy2 was found to be broadly expressed in V1R and V2R lineages, only sparse VSNs express Cnpy3 -4 and -5 (Fig. 2B).

**Figure 2.**
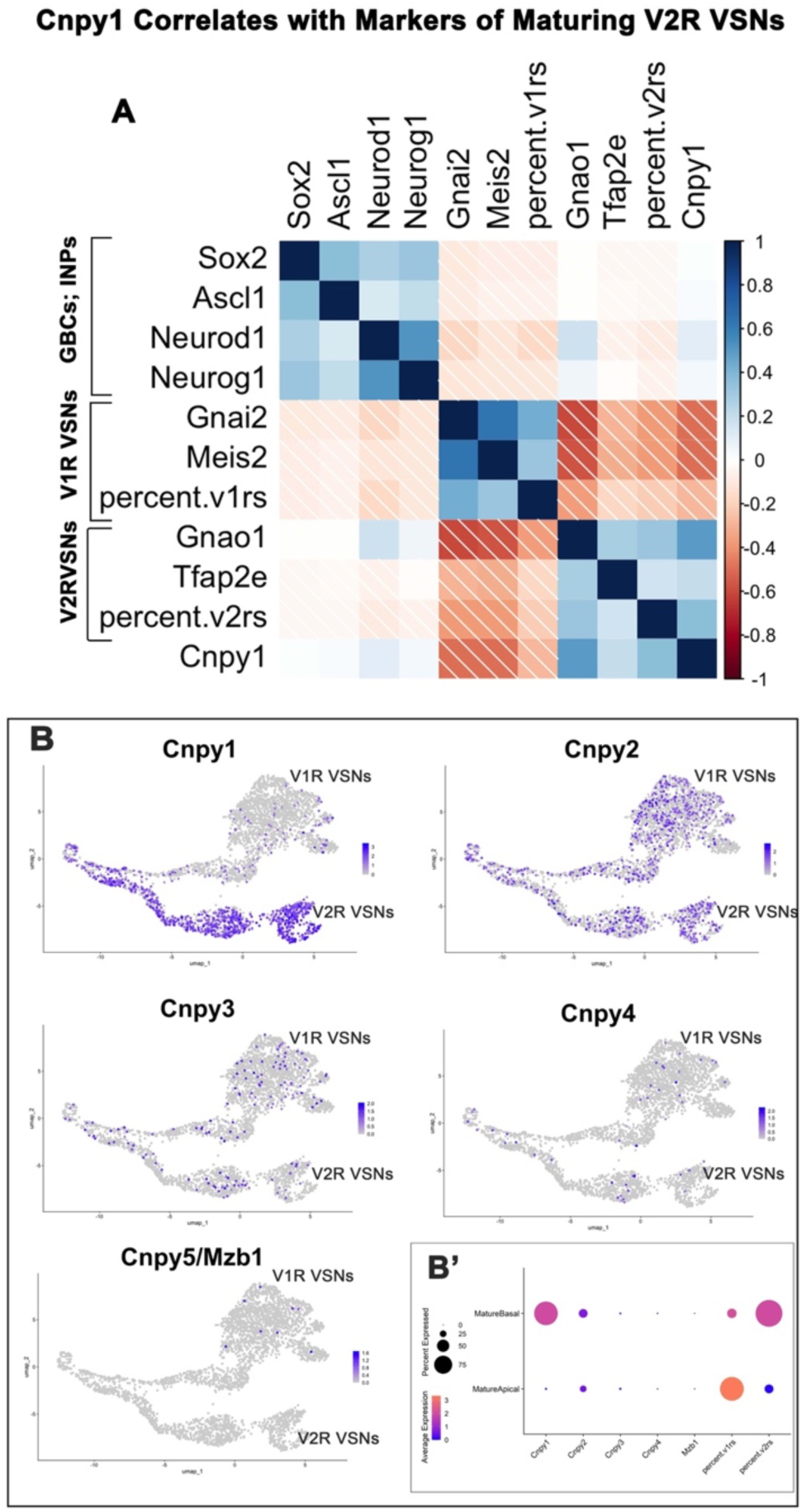
Cnpy1 expression correlates with the maturation program of basal V2R VSNs. (A) Correlation matrix showing that *Cnpy1* expression positively correlates with markers of differentiating and maturing basal VSNs, including *Gnao1* (Gαo), *Tfap2e*/AP-2ε, and the proportion of *V2R*-expressing neurons, and is inversely correlated with apical *V1R* markers. (B) Feature plots of *Cnpy1*, *Cnpy2*, *Cnpy3*, *Cnpy4*, and *Cnpy5/Mzb1* across the VSNs, UMAP show that only *Cnpy1* is selectively enriched in V2R VSNs. (B′) Dot-plot summarizing relative expression levels and percent-positive cells for CNPY family members in mature apical and basal VSNs, highlighting the selective basal enrichment of *Cnpy1*.

#### At two weeks after birth, despite the absence of Cnpy1, the VNO of Cnpy1KO has normal gross morphology

To explore the function of Cnpy1 in the VNO, we analyzed an unpublished Cnpy1 knockout mutant mouse line (C57BL/6J-Cnpy1^em1cyagen^) carrying a 6.2 kb deletion in the Cnpy1 coding sequence. Cnpy1 KOs are viable and fertile. Heterozygous mice were bred to produce Cnpy1^−/−^ mice and controls.

Immunolabeling against Cnpy1 and AP-2ε in P14 WT mice (Fig. 3) confirmed that Cnpy1 protein expression was limited to basal VSNs. In contrast, Cnpy1 KO littermates showed no immunodetectable Cnpy1 while still exhibiting AP-2ε immunoreactivity in the V2R VSNs. These data confirmed the absence of the protein in the KOs and the specificity of the commercially available Cnpy1 antibody (NBP2-82720 Novus Bio) (Fig. 3A, B). Furthermore, both major VSN subtypes were present in KO mice, with proper segregation and an overall morphology indistinguishable from WT controls two weeks after birth.

**Figure 3.**
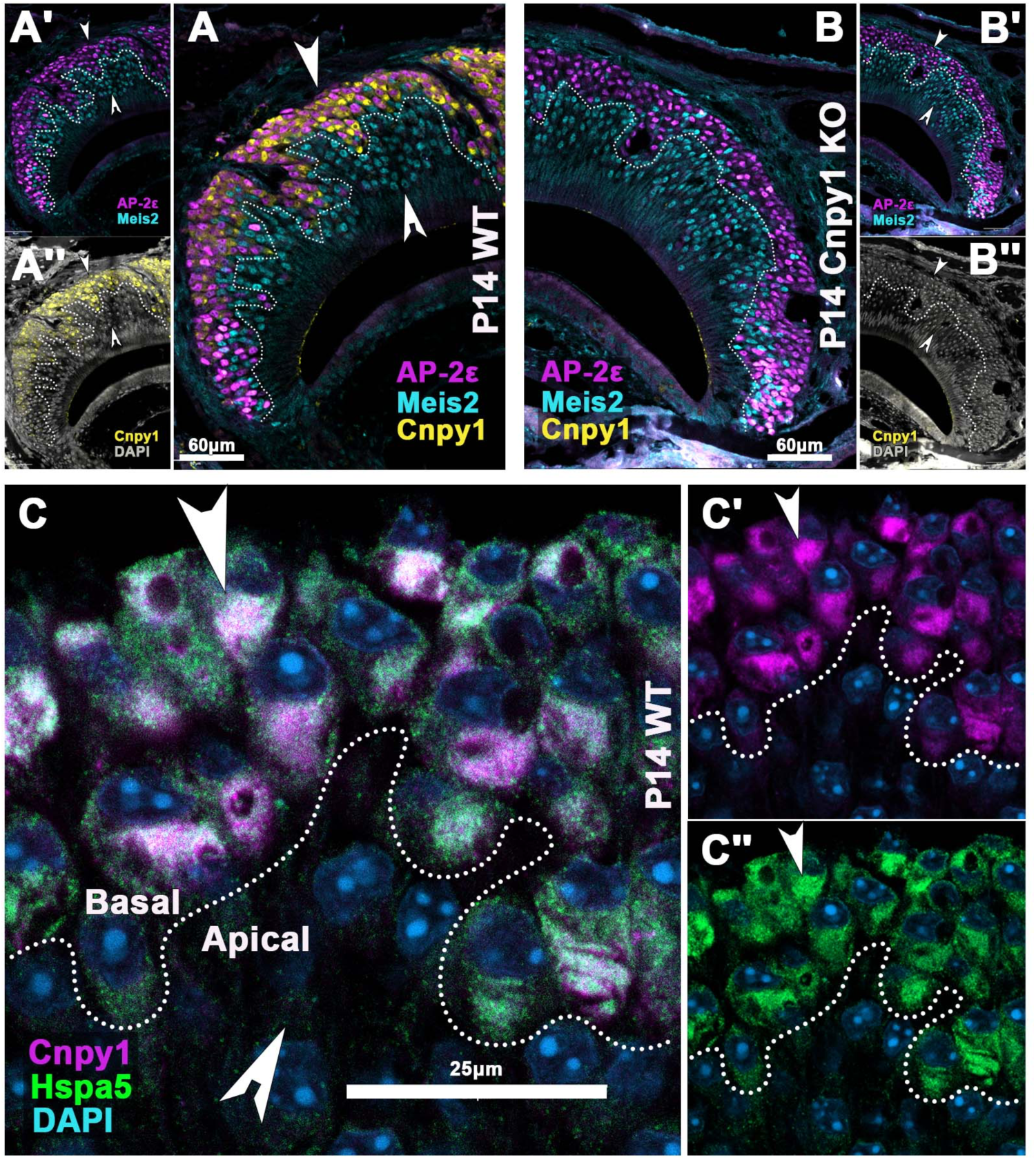
CNPY1 protein is enriched in basal VSNs and colocalizes with the ER chaperone HSPA5. (A–A′) Immunofluorescence for AP-2ε, Meis2, and CNPY1 in P14 WT VNO shows robust CNPY1 expression in basal V2R VSNs; a notched arrowhead marks apical-layer cells. (B–B′′) P14 *Cnpy1* KO VNOs show AP-2ε immunoreactivity, normal apical–basal layering but complete loss of CNPY1 immunoreactivity. (C–C′′) High-magnification images showing CNPY1 and the ER chaperone HSPA5 (BiP) colocalize in basal VSNs (arrowhead), whereas neither protein is detectable in apical VSNs (notched arrowhead). Scale bars: 60 µm (A–B), 25 µm (C).

Notably, different from the zebrafish Cnpy1 orthologue, and from the other members of the Cnpy1 family, the mammalian Cnpy1 lacks the ER localization signal (Schildknegt et al., 2019). Nonetheless, we observed overlapping expression patterns with the ER protein Hspa5, which suggests that Cnpy1 is an ER-resident protein in mammals (Fig. 3C).

#### Cnpy1 KO Mice Exhibit a Progressive Loss of V2R VSNs

We performed immunostaining against the transcription factors AP-2ε and Meis2 to quantify V1R and V2R VSNs at different developmental stages (Fig.4 A-E’). At P21, quantification revealed that the KO animals had a 13% decrease in the number of basal AP-2e expressing neurons (Fig. 4C). Examination of the VNOs at P60 (Fig. 4D-E’) showed a noticeable thinning of the basal VSN layer, amounting to a 42% reduction compared to the WT control (Fig. F). While apical VSN number was unchanged between P21 WT and KO mice, we observed a slight increase in P60 KO mice.

**Figure 4.**
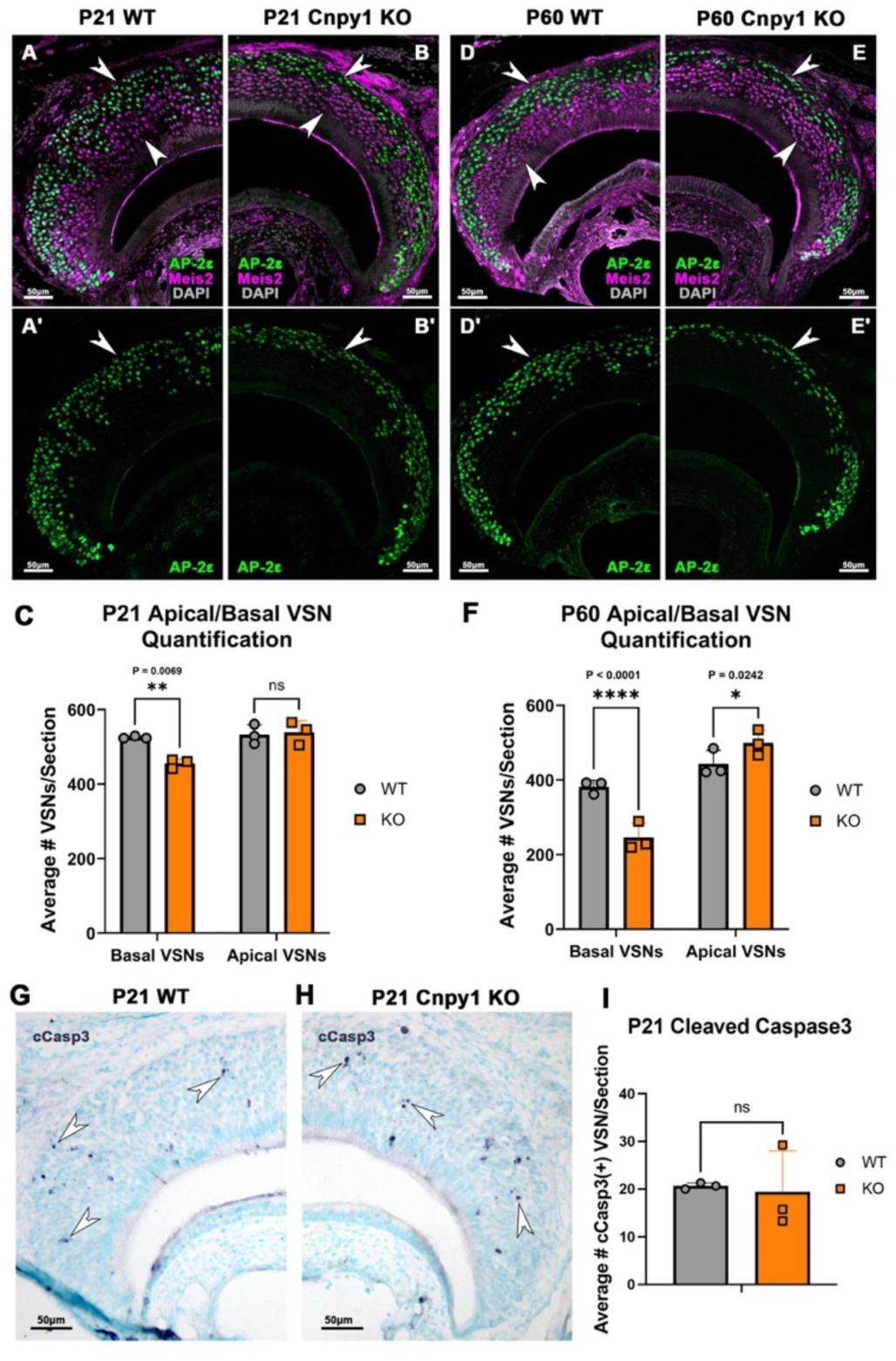
Loss of CNPY1 leads to progressive depletion of basal VSNs without a significant increase in apoptosis at P21. (A–A′) P21 WT VNO immunofluorescence for AP-2ε and Meis2 showing normal apical–basal organization. (B–B′) P21 *Cnpy1* KO sections display a mild but noticeable reduction in the AP-2ε⁺ basal VSN layer (arrowheads). (D–D′) At P60, WT tissue maintains a robust basal compartment. (E–E′) P60 *Cnpy1* KO VNOs show pronounced thinning of the AP-2ε⁺ basal layer and reduced Meis2⁺ V2R VSNs. (C, F) Quantification of AP-2ε⁺ basal and Meis2⁺ apical VSNs shows a significant reduction of basal VSNs in *Cnpy1* KO at P21 (p<0.0069) and a dramatic loss by P60 (p<0.0001), with apical VSN (P60 p<0.0242)numbers largely preserved. (G– H) Cleaved caspase-3 immunostaining in P21 WT and KO VNOs reveals comparable sparse apoptotic profiles (notched arrowheads). (I) Quantification confirms no significant increase (p > 0.05) in cleaved caspase-3⁺ VSNs in *Cnpy1* KO at P2, N=3, significance calculated unpaired two tailed t test analysis. Scale bars: 50 µm.

To assess whether the dramatic reduction between P21 and P60 could result from increased apoptosis, we performed immunohistochemistry for Cleaved Caspase-3 (cCasp3), a marker for apoptosis, at P21 and performed relative quantifications (Fig. 4C-I). Interestingly, as previously reported for other mouse lines (Gg8) (Montani et al., 2013), a reduction in basal was not associated with a significant difference in the number of Cleaved Caspase-3 positive VSNs found between controls and KO.

#### Transcriptomic analysis suggests a degenerative phenotype

scRNA sequencing of P21 WT and Cnpy1 KO mice was performed; the UMAP of WT and KO cells, represented in red and blue (Fig. 5A), respectively, shows a high degree of overlap, indicating a gross transcriptional similarity. The cells of the VNO were categorized by their markers of maturity and whether they fell along the apical or basal developmental path post-dichotomy (Fig 5B). For each group of cells along the basal developmental path, the number of significantly misregulated genes was defined based on (FDR <= 0.05). Notably, the number of differentially expressed genes increased with maturation stage (Fig. 5C), indicating a progressive neurodegenerative phenotype that begins prior to maturation of the neuron. We performed a GO term analysis for Neuronal Precursors, immature basal VSNs, and mature basal VSNs (Fig. 5E). A list of differentially expressed genes in mature basal VSNs is listed in Fig. 5E.

**Figure 5.**
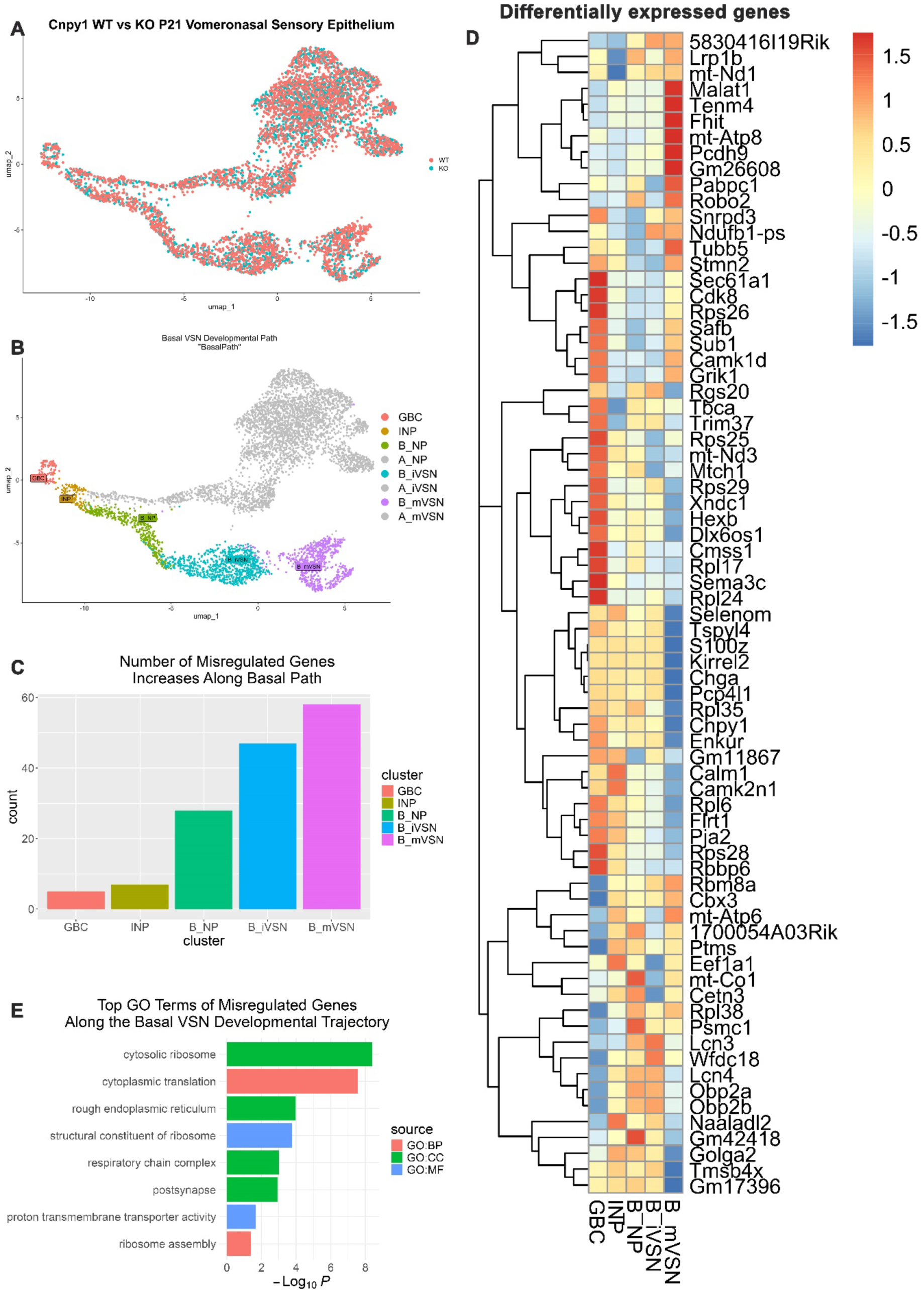
Single-cell transcriptomics reveals stage-progressive gene misregulation along the basal VSN developmental trajectory in *Cnpy1* KO mice. (A) UMAP overlay of WT (orange) and *Cnpy1* KO (teal) P21 VNO single-cell transcriptomes showing that both genotypes follow similar developmental trajectories from GBCs to mature VSNs. (B) Basal VSN differentiation path (“BasalPath”) subdivided into five transcriptional stages: GBC, INP, basal neuronal precursors (B_NP), basal immature VSNs (B_iVSN), and basal mature VSNs (B_mVSN). (C) Bar plot showing that the number of differentially expressed genes progressively increases as basal VSNs mature, with the greatest dysregulation occurring in B_mVSNs. (D) Heatmap of representative differentially expressed genes in B_mVSNs, highlighting broad transcriptional disruption in *Cnpy1* KO basal VSNs, (red) upregulated in Cnpy1KO; blue downregulated in the Cnpy1KO. (E) GO-term enrichment analysis of differentially expressed genes throughout the basal VSN trajectory, as shown by the simplified Driver Terms. Within these Enriched biological processes include pathways related to neurodegenerative diseases, protein folding and quality control, ribosome biogenesis, translation, and cellular homeostasis.

#### The progressive reduction of basal VSN is associated with a lack of chemo-sensory activity

Previous studies have shown that VSNs of mice carrying mutations in genes that impair VSN activation such as GαoKO, AP-2εKO, Smad4CKO, AtF5KO, Gγ8KO, TRPC2KO are progressively lost within two months after birth (Chamero et al., 2011; Lin et al., 2018; Montani et al., 2013; Naik et al., 2020; Nakano et al., 2016; Stowers et al., 2002). This prompted us to investigate whether the VSNs of Cnpy1 KO were functional (Fig. 6). To test VSN activity, male mice were exposed to male urine, which is known to activate both V1R and V2R VSNs. Immunofluorescent staining against phosphorylated ribosomal subunit S6 (pS6) (Silvotti et al., 2018) was performed as a proxy for VSN signal transduction (Fig. 6A-B’). The number of pS6+ basal VSNs was nearly absent in Cnpy1 KO mice (Fig. 6C), in stark contrast to WT controls, whereas the number of pS6+ apical VSNs was not significantly different from controls.

**Figure 6.**
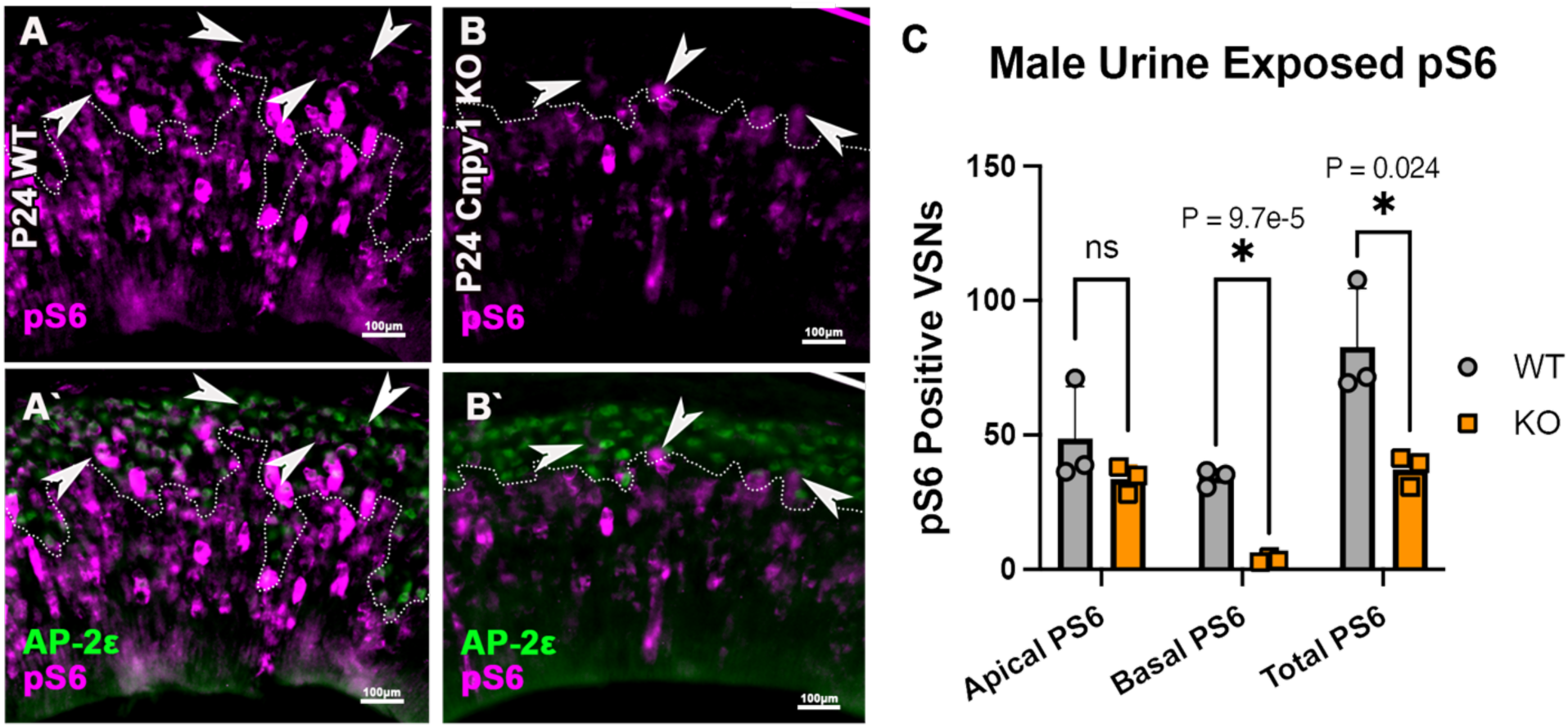
Basal VSNs fail to activate in response to male urine in *Cnpy1* KO mice. (A–A′) In P24 WT VNO, pS6 immunostaining reveals robust activation of both apical (AP-2ε⁻) and basal (AP-2ε⁺) VSNs following male-urine exposure (notched arrowheads). (B–B′) In *Cnpy1* KO mice, apical VSN activation is preserved, but activated pS6⁺ basal VSNs are nearly absent. (C) Quantification of pS6⁺ VSNs confirms a dramatic loss of activity in basal VSNs of *Cnpy1* KO animals (basal VSNs p<.00097, total PS6 p<.024), while apical responses remain comparable to WT, N=3, significance calculated BY log10 transformation on percent values; unpaired two tailed t test analysis. Scale bars: 100 μm.

#### V2R receptor expression levels are reduced in Cnpy1 KO VNOs, and V2Rs are not detected in the lumen of the VNO

Based on these results, we decided to check the expression of the receptors. Interestingly, scRNA sequencing data showed that Cnpy1 KO mice have increased V2R2 receptor gene expression (Fig. 7). To verify this, we performed immunofluorescence staining on P21 WT mice and Cnpy1 KO mice against AP-2ε, V2R2, and Hspa5. Hspa5, was used to highlight the ER of V2R neurons to create a three-dimensional mask of the ER using Imaris (Fig.7 A’’’, B’’’). V2R2 immunofluorescent intensity within Hspa5 regions was measured. To avoid bias from different sized regions of mask, total V2R2 signal was divided by total volume of Hspa5 mask to derive a density of V2R2 immunofluorescence. Cnpy1 KO mice showed a significant reduction of V2R2 signal density (Fig. 7C). Despite higher mRNA expression in Cnpy1 KO mice, lower detectability of the protein is observed suggestive of reduced translation or defective protein processing.

**Figure 7.**
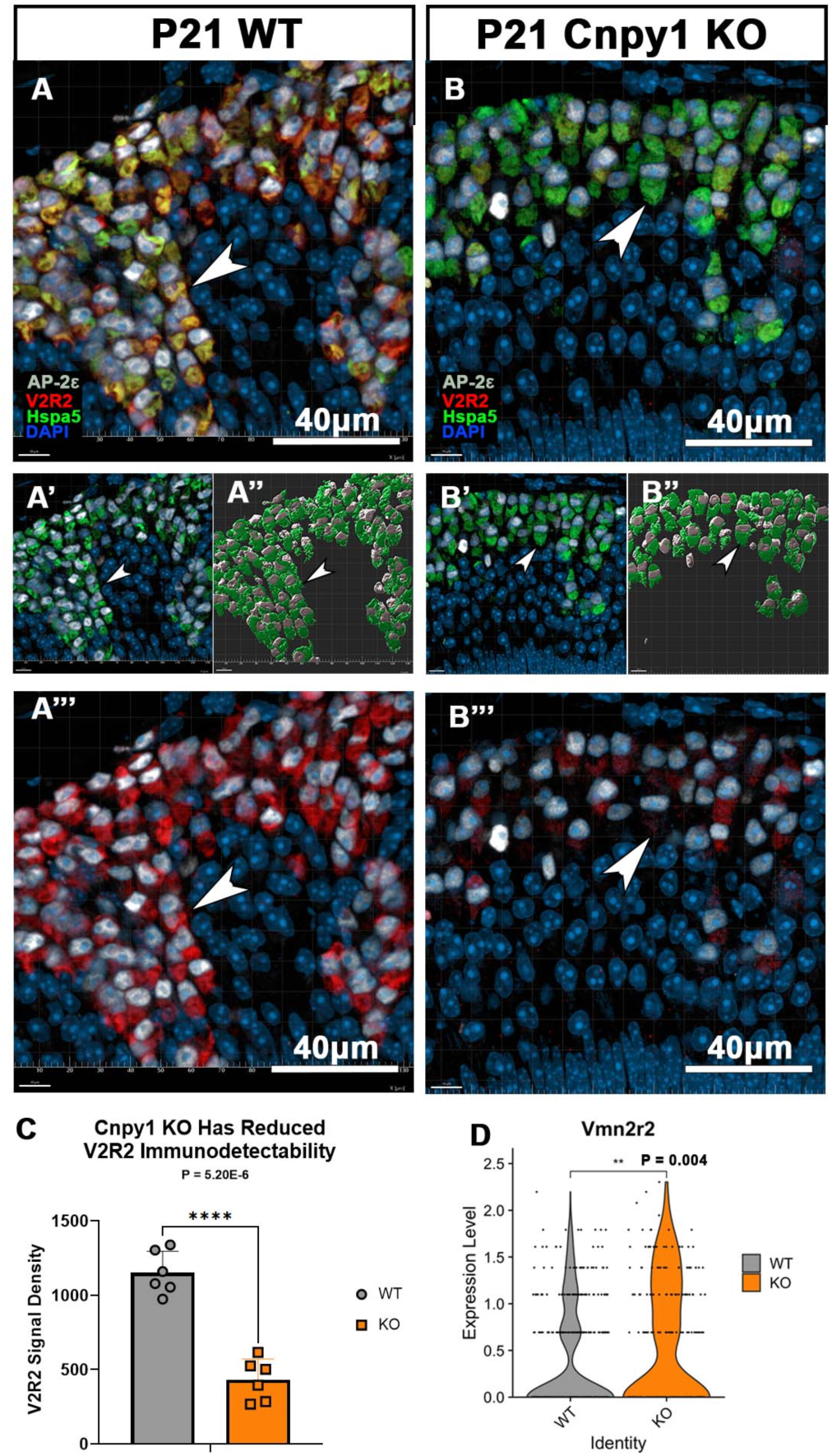
CNPY1 loss reduces V2R2 protein detectability despite increased transcript levels. (A–A‴) P21 WT VNO stained for AP-2ε, V2R2, HSPA5, and DAPI. HSPA5 marks the ER of basal V2R VSNs and was used to generate 3D ER masks in Imaris for quantification (arrowheads). (B–B‴) P21 *Cnpy1* KO sections show markedly reduced V2R2 immunofluorescence within HSPA5-defined ER regions despite intact ER structure. (C) Quantification of V2R2 signal normalized to total HSPA5-mask volume reveals >60% reduction in V2R2 protein density in *Cnpy1* KO mice (p<0.000052) N=3, significance calculated by log10 transformation on percent values; unpaired two tailed t test analysis. (D) scRNA-seq analysis shows significantly increased *Vmn2r2* mRNA levels in *Cnpy1* KO basal VSNs (p<0.004, using stat_means, indicating discordance between transcript abundance and protein detectability. Scale bars: 40 µm.

We applied a similar masking strategy to assess whether V2R receptors were correctly transported to the lumen, the site of external signal detection (Fig.8). WT and Cnpy1 KO VNOs at P21 were stained for V2R2 and Gαo (Fig. 8). The Gαo signal was used to generate a three-dimensional mask delineating the lumen (Fig. 8A′, B′), and V2R2 signal intensity within this masked region was quantified and normalized to the mask volume (Fig.8). In WT mice, clear V2R2 immunoreactivity was detected at the lumen (arrows), whereas Cnpy1 KO mice exhibited a near-complete loss of detectable V2R2 signal in this region.

**Figure 8.**
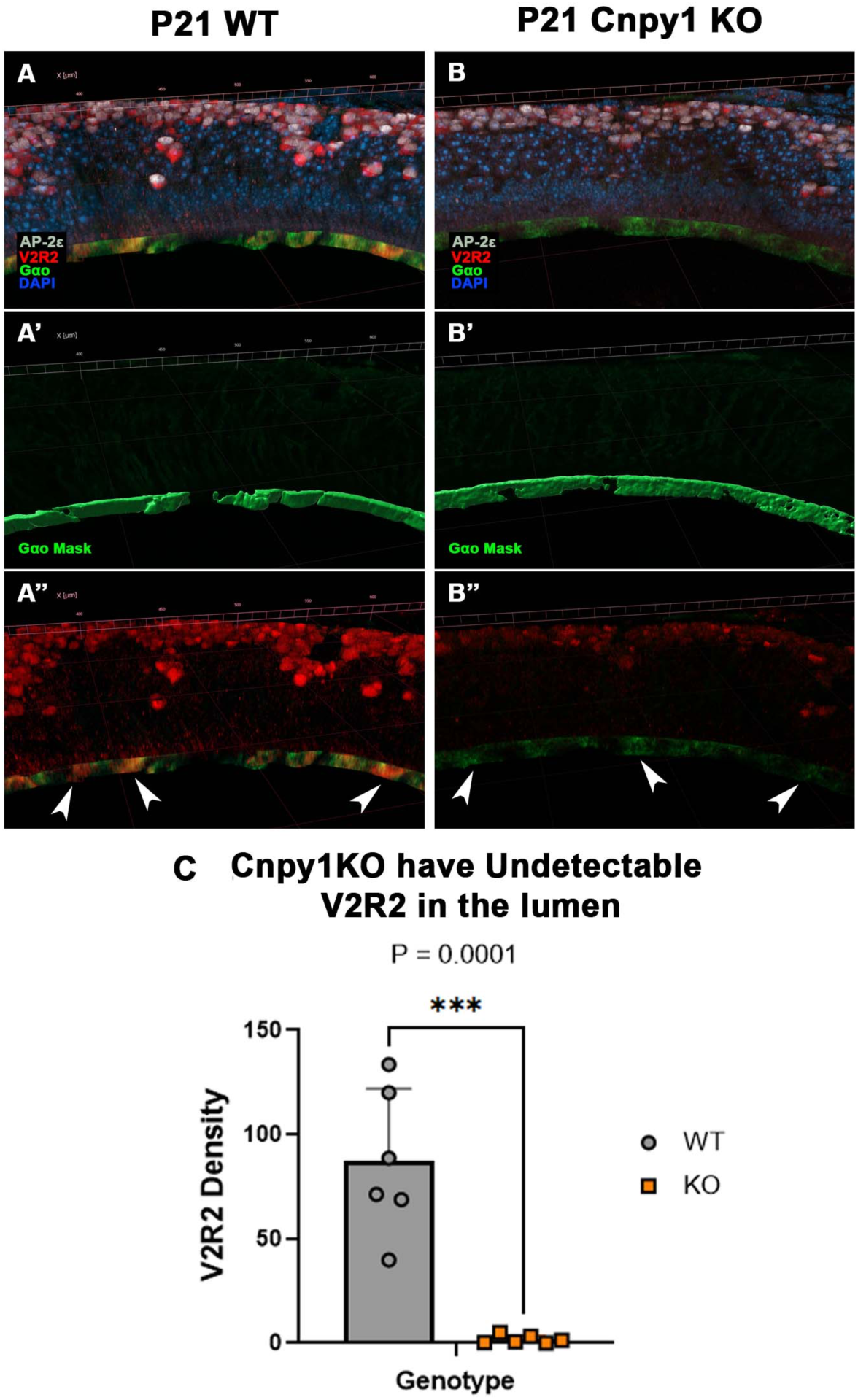
CNPY1 is required for proper trafficking of V2R2 to the luminal surface of basal VSNs. (A–A″) P21 WT VNO stained for AP-2ε, V2R2, Gαo, and DAPI shows robust V2R2 immunoreactivity at the luminal surface (arrows). Gαo signal was used to generate a 3D mask defining the lumen for quantitative analysis. (B–B″) In *Cnpy1* KO mice, V2R2 signal within the Gαo-defined luminal mask is nearly absent despite preserved Gαo labeling. (C) Quantification of V2R2 density normalized to luminal mask volume reveals a near-complete loss ((p<0.0001) N=6, significance calculated log10 transformation on percent values; unpaired two tailed t test analysis) of luminal V2R2 in *Cnpy1* KO mice. Scale bars: 40 µm.

#### ER stress gene expression changes during maturation, and Cnpy1 KO shows signs of increased ER stress

Recent findings suggest that ER stress levels in chemosensory neurons are driven by the expression of odorant receptors and that ER stress can directly regulate the expression of axon guidance and adhesion molecules, including Robo2, Class III Semaphorins, Kirrels, and protocadherins, as part of a stress-responsive transcriptional program (Shayya et al., 2022). Notably, the expression of some of these molecules, such as Kirrel2, are also regulated by neuronal activity.

In Cnpy1 knockout mice, we observed markedly reduced immunoreactivity for V2R receptors, consistent with defects in receptor processing and trafficking. Impaired ER processing is likely to increase ER stress and activate compensatory unfolded protein response mechanisms that can ultimately alter global protein translation.

To determine if defective V2R processing could be reflected by changes in ER stress, we analyzed the expression of genes activated by ER stress compensatory mechanisms, including canonical players such as Atf6, Ddit3 (CHOP), Hspa5 (BiP), and Xbp1(Remondelli and Renna, 2017), and the activating transcription factor 5 (Atf5)(Wang et al., 2023) (Fig. 9A). This analysis revealed 1) that ER stress increases from immature VSN to mature (Fig 9A); in fact, Xbp1, Atf6, Ddit3/CHOP, Selenos, Manf, and Hspa5 levels increased in both WT and Cnpy1 KOs as the neurons matured. 2) ER stress elevation coincides with V2R expression. In fact, immunohistochemistry and transcriptomic data revealed that Ddit3/CHOP expression increases as the VSNs express V2R (Fig.9 B-G). 3) In the KO, we found increases in mRNA levels of Xbp1, Atf5, Ddit3/CHOP, and Hspa5 in mature neurons, which suggest increases in intracellular stress (Fig. 9A). Notably, ER stress and compensatory mechanisms can alter translation (Ron, 2002; Wolzak et al., 2022). Supporting this, analysis of transcriptional data from WT and Cnpy1 KO revealed variations in the expression of multiple genes of the Ribosomal Cellular Component (GO:0005840) (Fig. 9H).

**Figure 9.**
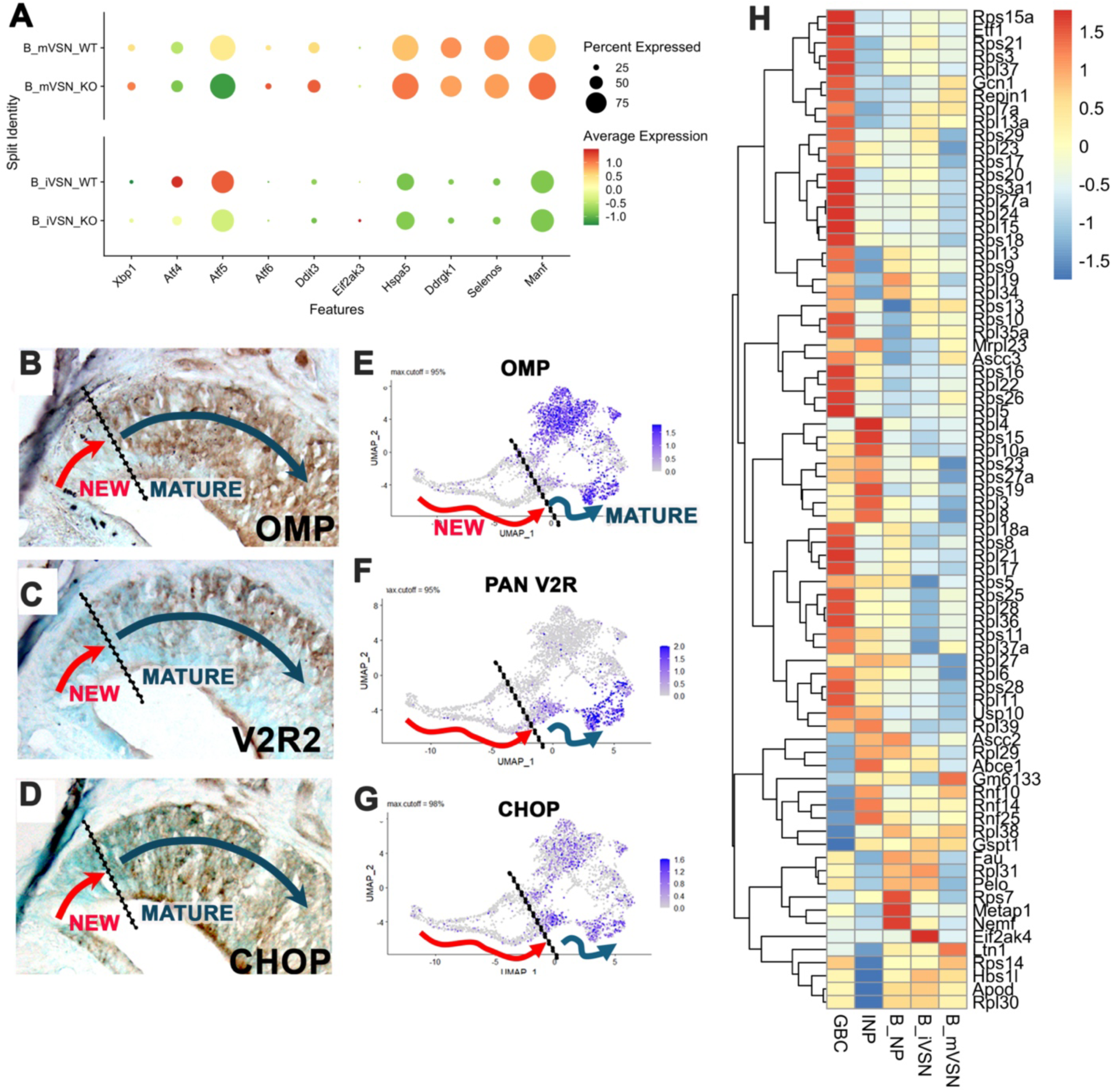
ER stress gene expression and maturation-dependent changes in vomeronasal sensory neurons (VSNs). (A) Dot-plot showing expression of key ER stress/UPR-related genes across immature (iVSN) and mature (mVSN) V2R-expressing neurons in WT and Cnpy1 KO mice. Classic UPR markers, including Atf6, Ddit3 (CHOP), and Hspa5 (BiP), are expressed at low levels in immature neurons and increase as VSNs mature, in both genotypes. Mature VSNs exhibit stronger ER-stress signatures. (B–D) Immunohistochemistry on P15 VNO sections showing spatial maturation of VSNs in the marginal zone. Immunoreactivity for OMP (B), V2R2 (C), and CHOP (D) increases along the gradient from newly generated neurons (NEW) toward fully mature cells (MATURE), indicating coordinated upregulation of sensory receptors, maturation markers, and ER-stress/UPR effectors. (E–G) Feature plots from scRNA-seq showing the distribution of OMP (E), pan-V2R expression (F), and CHOP/Ddit3 (G) along the developmental trajectory. Gene expression intensity mirrors the histological maturation gradient, with all three markers increasing as VSNs transition into the mature V2R-expressing state. (H) Heatmap displaying differential expression of ribosomal protein genes (GO:0005840 Ribosome Cell Component Aspect), ribosome-associated factors, and translation-related genes across GBCs, iNPs, NP, immature VSNs, and mature VSNs in WT versus Cnpy1 KO mice. The analysis highlights genotype-dependent alterations in components of the translational machinery that accompany disrupted ER proteostasis in Cnpy1-deficient VSNs. Red represents higher expression levels in the WT and blue in the Cnpy1 KO.

#### Cnpy1 KO shows changes in adhesion and guidance molecules and disorganized glomeruli in the AOB

Variation in ER stress can alter the expression of genes involved in axonal guidance and coalescence (Shayya et al., 2022). Strikingly, single-cell analysis (Fig. 10A) revealed a near-complete loss of Sema3C mRNA and clear changes in the expression of multiple other guidance genes, including Epha5, Kirrel2, Pcdh9, Pcdh17, and Robo2, in mature V2R-VSs of the Cnpy1KOs.

**Figure 10.**
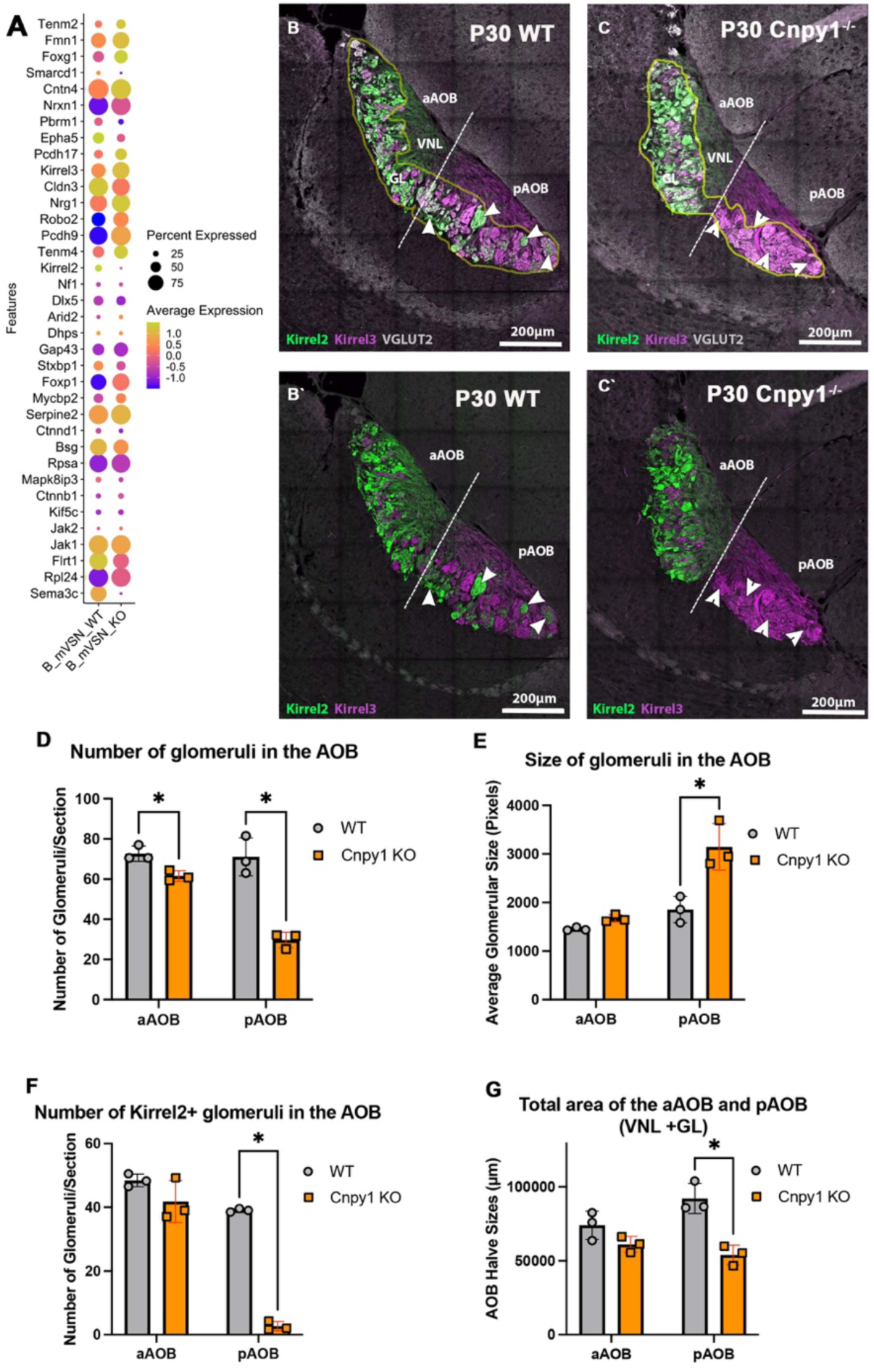
Loss of CNPY1 alters the expression of adhesion and guidance molecules and disrupts posterior AOB organization and glomerular architecture. (A) Dot plot showing the expression of Cell Adhesion molecules and Axonal Guidance genes in the V2R/VSNs of WT and Cnpy1KOs. (GO:0007155 and GO:0007411 respectively) (B–B′) P30 WT AOB sections stained for Kirrel2 (green), Kirrel3 (magenta), and VGLUT2 (grey) illustrate normal segregation of aAOB and pAOB glomeruli. (C–C′) P30 *Cnpy1*⁻/⁻ sections show reduced pAOB territory and aberrant Kirrel2⁺/Kirrel3⁺ glomerular patterning. (D) Total VGLUT2⁺ glomeruli in the AOB are decreased in *Cnpy1*⁻/⁻ mice(aAOB p<0.013294, pAOB p<0.002169). (E) Average size of VGLUT2⁺ glomeruli are significantly increased in the pAOB of Kos (p<0.000463), indicating defective glomerular compaction. (F) The number of Kirrel2⁺/VGLUT2⁺ pAOB glomeruli is reduced in KOs (p<0.000002) (G) Quantification of aAOB and pAOB (vomeronasal nerve layer + glomerular layer) total area shows a significant reduction in the pAOB (p<0.005581) of *Cnpy1* KO mice N=3, significance calculated unpaired two tailed t test analysis. Scale bars: 200 µm.

Previous studies showed that Kirrel2 expression is reduced in Trpc2 knockout mice, which lack pheromone-evoked activity (Stowers et al., 2002), demonstrating that Kirrel expression is activity-dependent in VSNs (Prince et al., 2013). Consistent with our transcriptomic data and with the loss of V2R-mediated sensory activity in Cnpy1 knockouts, immunostaining revealed an almost complete loss of Kirrel2 in the posterior accessory olfactory bulb, accompanied by an abnormal expansion of Kirrel3 expression across all glomeruli. Kirrel proteins regulate the homophilic adhesion and coalescence of like-with-like olfactory axons, thereby controlling the proper size and number of glomeruli in the olfactory map (Prince et al., 2013). Quantification of glomeruli number and size showed an overall reduction of the posterior AOB (Fig. 10G), which, like for other mutants, reflects a reduction of basal VSNs. However, we observe a much more dramatic reduction in the number of glomeruli, close to 50%, together with increased glomerular size (Fig. 10D, E) in Cnpy1 KO compared to controls. These phenotypes largely resemble those of Kirrel mutants (Prince et al., 2013).

#### AP-2εKO mice have reduced expression of Cnpy1

Previous studies show that the identity of V2R-expressing VSNs is established by early Notch activation, followed by increased Bcl11b and subsequent induction of AP-2ε (Enomoto et al., 2011; Katreddi et al., 2022; Lin et al., 2022). This transcriptional cascade coordinates the expression of the molecular machinery required for basal-cell specialization, likely including the ER-resident proteins and chaperones needed for proper V2R processing and trafficking. AP-2ε has been proposed to upregulate Calr-4 and Cnpy1, both of which appear to directly interact with V2R proteins and likely control their proteostasis (Devakinandan et al., 2025; Dey and Matsunami, 2011). The degenerative phenotype of Cnpy1 KO mice partially overlaps with that of AP-2ε KO mutants (Lin et al., 2018). We therefore examined Cnpy1 expression in AP-2ε KO mice at P10, before the epithelium gets severely disrupted.

For this analysis, we immunolabeled vomeronasal sections from WT and AP-2ε KO mice at against Meis2, Hspa5, and Cnpy1 (Fig. 11). This staining allowed us to identify the basal VSNs based on the lack of Meis2 expression; Hspa5 was used to generate masks for the presumptive ER, and the signal intensity was measured. The results indicate that AP-2ε KO mice have a significant reduction in Cnpy1 expression.

**Figure 11.**
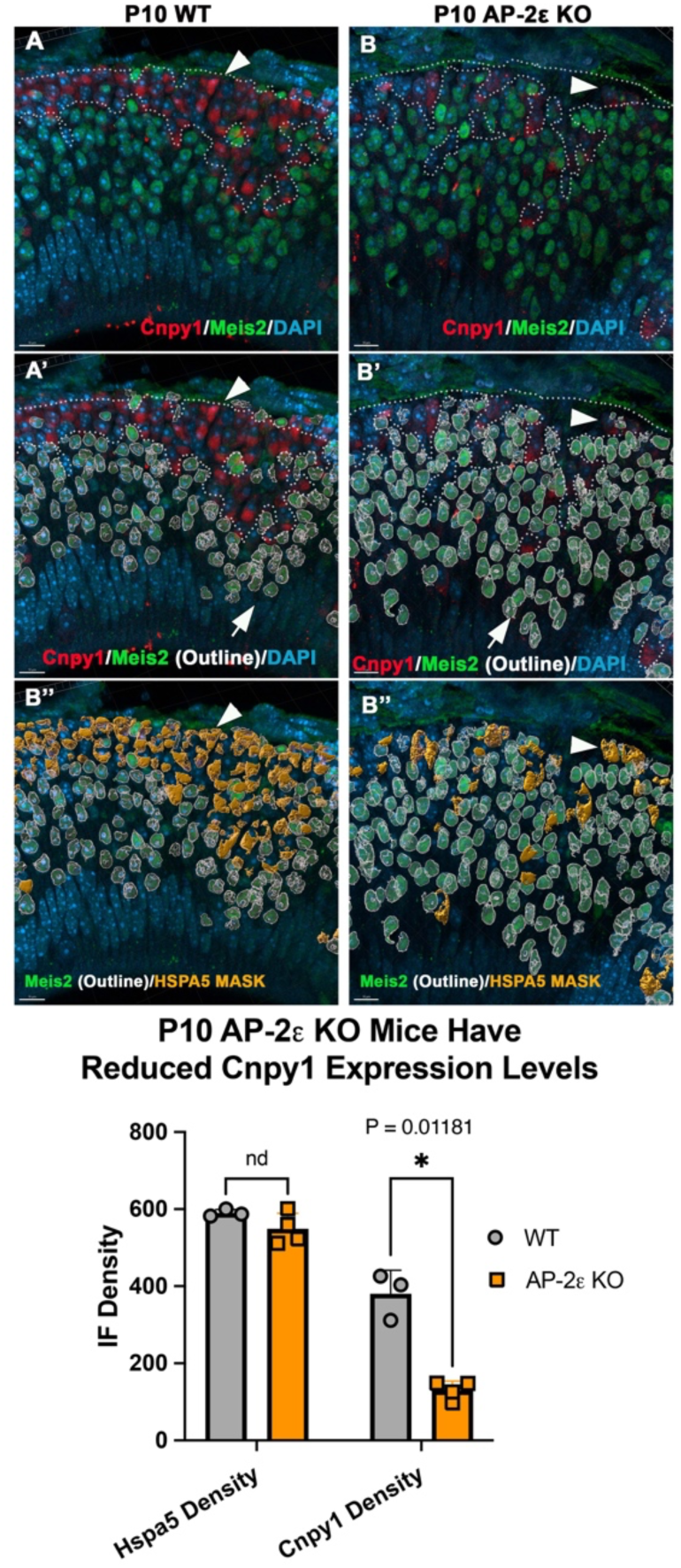
AP-2εKOs have reduced Cnpy1 expression in basal V2R VSNs. (A–B) Representative P10 WT and AP-2ε KO VNO sections stained for Cnpy1 (red; arrowheads), Meis2 (green), and DAPI (blue). Dashed lines indicate apical and basal territories, which are clearly defined in WT but less distinct in KO samples. Meis2-negative cells were classified as basal VSNs. (A′–B′) Outline masks highlight Meis2-positive apical VSNs. In WT samples, Cnpy1 is strongly expressed in basal V2R-expressing VSNs, whereas AP-2ε KOs show markedly reduced Cnpy1 immunoreactivity. (A″–B″) Hspa5 immunostaining was used to generate 3D ER-based masks of basal VSNs for quantitative analysis. (C) Quantification of Cnpy1 immunofluorescence density within Hspa5-defined basal VSN masks. AP-2ε KO mice show a significant reduction in Cnpy1 levels compared to WT controls, while Hspa5 density is unchanged. Data are presented as mean ± SEM.

#### Loss of Fgfr1 in newly formed VSNs does Not Recapitulate Cnpy1 KO Degenerative Phenotype

Prior to our study and another similar study, Cnpy1 has been characterized only in zebrafish where it was proposed to be a key regulator for FGFR1 expression (Matsui et al., 2011). While Cnpy1 is expressed throughout the development of the V2R VSNs, FGFR1 is transiently expressed in newly formed VSNs. (Fig. 12A-A’). To test whether Cnpy1 phenotypes could partially result from defective FGFR1 signaling, we crossbred Fgfr1^fl:/fl:^ mice with Neurogenin1-CreERt2 /Fgfr1 ^fl:/fl:^/A14^+/+^ reporter mice. We previously used this approach (Katreddi et al., 2022) to induce chimeric recombination in newly formed VSNs. Tamoxifen injections at P7 induced Cre-mediated recombination in newly generated apical and basal VSNs, which was monitored by the Ai14Rosa-tdTomato reporter. Mice were analyzed at P60 (53 days post-injection). Notably, we observed no significant differences between Ngn1-CreERt2^+/-^;Ai14^+/-^ and Ngn1-CreERt2^+/-^;Ai14^+/-^;Fgfr1^fl:/fl:^ mice. The proportion of tdTomato-labeled VSNs co-expressing either AP-2ε or Meis2 was comparable across genotypes, indicating that loss of Fgfr1 does not lead to VSN degeneration, in contrast to the pronounced neuronal loss observed in the Cnpy1 KO phenotype (Fig. 12).

**Figure 12.**
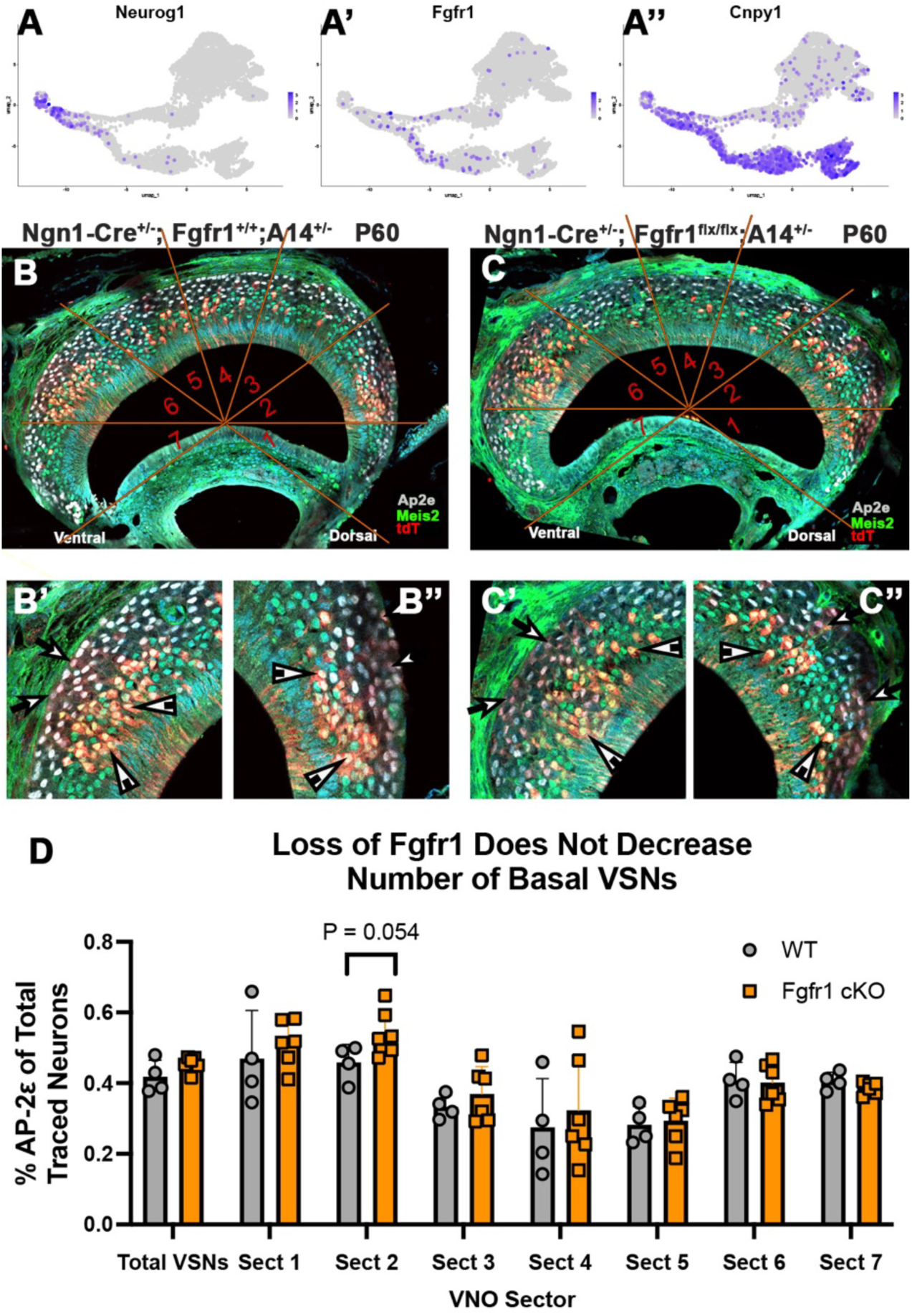
FGFR1 loss does not phenocopy the *Cnpy1* KO and does not cause basal VSN loss. (A) Feature plots showing *Neurod1* (immature VSN marker), *Fgfr1*, and *Cnpy1* expression across the VSN UMAP. *Fgfr1*is expressed in immature neurons but does not overlap with the basal ER-associated program marked by *Cnpy1*, indicating distinct functional pathways.(B–B′′) *Ngn1*-CreERT2; *Fgfr1*⁺/⁺; *Ai14*⁺/⁻ control mice at P60 (53 DPI) show robust recombination in both apical (Meis2⁺) and basal (AP-2ε⁺) VSNs (arrowheads). Basal neurons exhibit lower tdTomato intensity due to reduced *Rosa2C*expression basal lineage-wide. (C–C′′) *Ngn1*-CreERT2; *Fgfr1*^flx/flx; *Ai14*⁺/⁻ conditional mutants show a similar pattern of recombination in apical and basal VSNs, with no evidence of basal VSN depletion. Recombined neurons are also present in the juvenile organ (JO). (D) Quantification across VNO sectors shows no significant reduction in the proportion of AP-2ε⁺ (basal) VSNs in *Fgfr1*cKO compared with controls, except for a small increase in one sector (Sect 2 p<0.054), WT N=4, Fgrf1 cKO N=6,significance calculated log10 transformation on percent values; unpaired two tailed t test analysis. Overall, basal VSN numbers remain stable, indicating that FGFR1 loss does **not** mimic the pronounced basal neuron loss observed in *Cnpy1* KO mice.

## Discussion

The vomeronasal organ (VNO) shows how evolution shapes neural diversity. Over millions of years, rodents have evolved a complex sensory epithelium that enables them to detect a wide range of social and environmental cues. The mouse VNO is among the most complex, with two main types of sensory neurons V1R- and V2R-expressing neurons and hundreds of subtypes. To better understand the molecular distinctions between VSN cell types, we utilized single-cell transcriptomics and mouse genetics.

Consistent with recent reports, we also found that V1R and V2R-expressing neurons differ by approximately 980 genes. Strikingly, the largest functional category of differentially expressed genes encodes proteins involved in endoplasmic reticulum (ER) synthesis, function, and protein processing. This observation highlights an underappreciated layer of neuronal diversity: beyond transcriptional regulation, the ER is the organelle responsible for folding, modifying, and trafficking proteins. The ER may itself define neuronal diversity by establishing the competence of neurons to produce certain sets of proteins within the constraints of its transcriptome. If the composition of ER-associated proteins differs among neuronal types, do all neurons manage proteostasis and ER stress similarly? Could cell type differences in ER composition/repertoire determine the idiosyncratic vulnerability of neurons to stress, misfolding, or neurodegeneration?

Within the basal VSNs, which express V2R receptors, we and others identified distinct ER protein signatures compared to apical VSNs. Morphological analyses further suggest structural differences in the ER between these neuronal populations. *Cnpy1 is* the among most highly expressed genes in V2R VSNs and is the most significantly differentially expressed ER-associated gene. Among the various differentially expressed ER proteins, we also observed that HSPA5/BiP, a key regulator of ER homeostasis involved in protein folding and modulation of stress response (Daverkausen-Fischer et al., 2022), is highly enriched in V2R VSNs within the VNO. This differential HSPA5/BiP expression suggests that V2R neurons may have distinct basal ER stress levels and varying capacities to manage ER stress compared to V1R neurons.

*Cnpy1* encodes a member of the evolutionarily conserved saposin-like protein family (Cnpy1– Cnpy4 in mammals). Cnpy proteins have been identified in species ranging from *Drosophila* and *C. elegans* to zebrafish, chick, and mammals. In the VNO we observed broad Cnpy2 expression. Apart from the findings presented here and those reported in a simultaneous independent preprint (Devakinandan et al., 2025), *Cnpy1* has previously been studied only in zebrafish at the midbrain– hindbrain boundary and in cell culture was found to function as a chaperone influencing FGFR1 maturation and FGF8 signaling. Rodent Cnpy1 does not have the ER retention signal; however, immunostaining against the ER marker Hspa5 and Cnpy1 suggests its expression in the ER. Notably, our analysis of FGFr1 conditional KO does not reveal overlap between Cnpy1 LOF and defective FGF signaling in the VNO (Fig. 12), suggesting that in the VNO Cnpy1 may have other functions or that compensatory mechanisms may mask the loss of Fgfr1.

In the VNO, Cnpy1 mRNA is expressed from globose basal cells through mature V2R-expressing VSNs (Fig.2). Morphological and transcriptomic analyses of control and Cnpy1 KO mice indicate that Cnpy1 loss does not prevent VSN neurogenesis or maturation. However, some genetic changes begin before maturation (Fig.5). Consistent with this, in V2R VSNs the phenotype emerges progressively: quantitative analyses at P21 and P60 show a gradual loss of basal VSNs (13% at P21; 42% at P60), indicative of a degenerative process. V1R/apical neuron numbers were unchanged at P21 and slightly increased at P60, possibly reflecting tissue redistribution or compensatory proliferation secondary to V2R loss. Notably small changes were also observed in V1R VSN connectivity, suggesting that defective Cnpy1 in newly formed neurons can have effects later (Fig.10). An alternative interpretation is that the alterations detected in the aAOB may arise secondarily from defects originating in the pAOB.

Cleaved Caspase-3 staining at P21 did not show apoptosis levels proportional to the extent of cell loss, indicating that neurodegeneration occurs with limited detectable apoptosis, similar to what was reported in Gγ8 mutant mice (Montani et al., 2013). However, we cannot rule out increased apoptosis at earlier or later stages.

Functional testing using male-urine stimulation and pS6 immunolabeling (Silvotti et al., 2018) showed that V2R-expressing neurons in Cnpy1 KO mice fail to activate. This prompted us to examine receptor expression and trafficking. V2R2, the most broadly expressed Family C receptor with reliable antibodies (Silvotti et al., 2011), showed markedly reduced somatic signal and an almost complete absence of protein at the dendritic lumen. Strikingly, these protein-level defects contrasted with the increased V2R2 mRNA detected in Cnpy1 KO scRNA-seq data, suggesting impaired translation, processing, and/or trafficking of V2R2 in the absence of Cnpy1. Similar to what has been proposed for Calr4, these data, together with what is proposed by others, suggest that Cnpy1 is a key component for V2R processing or trafficking (Devakinandan et al., 2025; Dey and Matsunami, 2011). However, single-cell data indicate altered expression of elongation factors and ribosome-associated proteins. At this stage, we cannot determine whether these changes directly contribute to the reduced V2R protein levels or whether they arise secondarily from increased intracellular stress. Transcriptomic analyses of the Cnpy1 KO also revealed that ER stress physiologically increases in parallel with V2R expression (Fig.9) and that, in Cnpy1 KO, UPR-related genes are upregulated compared to controls, suggesting increased ER stress levels.

A recent study proposed that the intrinsic sequence variability among odorant receptor genes can elicit differential levels of ER stress and stress responses within sensory neurons, and that these distinct ER stress signatures, in turn, regulate the expression of adhesion molecules and axon guidance cues (Shayya et al., 2022). This indirectly implies that the ER’s molecular repertoire and the efficiency of protein processing can determine neuronal wiring by altering adhesion dynamics and connectivity. Consistent with this model, our analysis of basal VSN connectivity in Cnpy1 knockout mice revealed disorganization of glomerular architecture in the posterior accessory olfactory bulb (AOB), including marked alterations in glomerular number and size, associated with stark changes in guidance and adhesion molecule expression. However, the basal VSNs of the Cnpy1 KO are nonfunctional, and previous studies have demonstrated that neuronal activity can also modulate the expression of key adhesion molecules; thus, in sensory systems, disruptions in signal transduction can indirectly reshape circuit assembly. In line with this observation, the pAOB of Cnpy1 KO mice exhibited a near-complete loss in expression of Kirrel2. Kirrel2 expression has been shown to depend on neuronal activity (Prince et al., 2013). However, distinguishing whether guidance and adhesion gene transcriptional alterations arise primarily from the loss of neuronal activity, from disrupted ER homeostasis, or from both remains challenging. We believe that this limitation applies not only to our study but also more broadly to efforts aimed at disentangling the relative contributions of proteostatic stress and activity-dependent transcriptional programs in shaping neuronal identity and connectivity.

Basal VSN identity is established through sequential activation of Notch signaling, Bcl11b, and ultimately AP-2ε, which drives the basal-specific transcriptome. Ectopic AP-2ε expression in apical neurons demonstrated its role as a master regulator, inducing ∼30% of basal-enriched genes, including the ER proteins Cnpy1 and Cal4, while repressing apical-enriched genes, including Calr and Nsg1 (Lin et al., 2022). These findings highlight that the ER-protein repertoire is an integral component of the basal gene regulatory network governed by AP-2ε, with Cnpy1 emerging as a key factor required for maintaining V2R proteostasis (see model in Fig.13). Our new data from AP-2e KO, where we observe reduced Cnpy1 expression, support this model (Fig. 11).

**Fig. 13.**
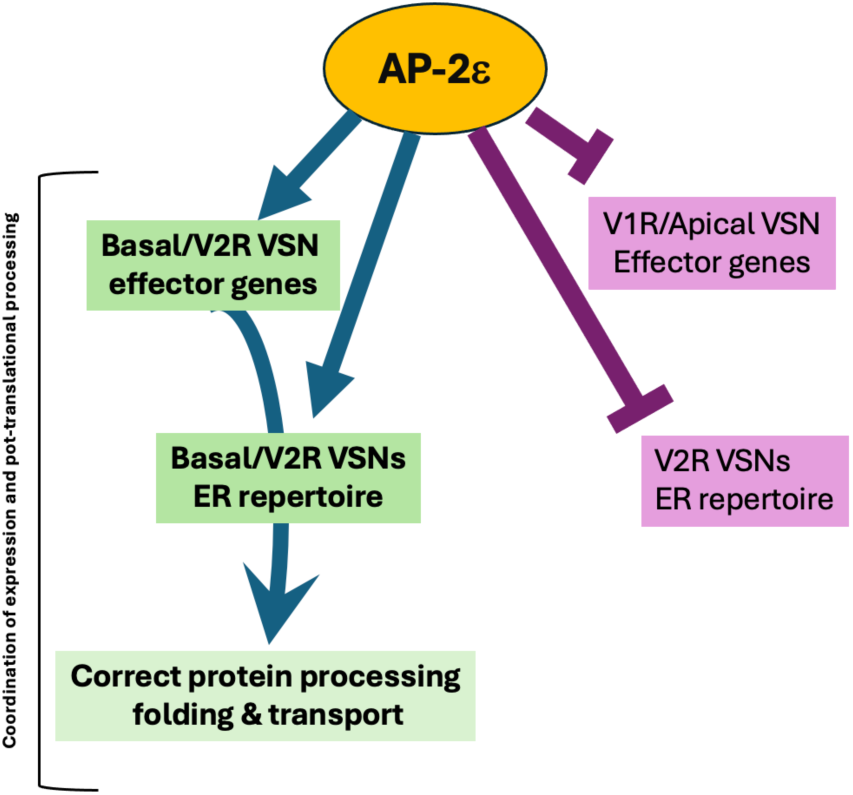
Model summary. AP-2ε acts as a master regulator of the basal/V2R VSN lineage by directly activating basal effector genes and the specialized ER proteostasis repertoire required for proper V2R protein folding, processing, and trafficking. At the same time, AP-2ε represses apical/V1R effector genes and apical-type ER components, thereby enforcing lineage specificity. This coordinated activation and repression ensures that basal VSNs acquire the correct molecular environment to process V2R receptors and establish functional connectivity.

While this manuscript was being prepared, an independent study (Devakinandan et al., 2025) reached many similar conclusions to those here presented using a different Cnpy1KO mouse. These independent results reinforce the finding that Cnpy1 is essential for proper V2R expression and function. Together, these studies underscore the importance of cell–type–specific ER chaperone networks in shaping receptor proteostasis and point to the broader value of systematically analyzing neuron-specific ER repertoires to understand both normal development and mechanisms of neurodegeneration.

## Materials and Methods

### Animals: Cnpy1KO(C57BL/6J-Cnpy1^em1cyagen^) mouse line

This Mouse line was generated for us by Cyagen Biosciences using CRISPR/Cas9-mediated deletion. Briefly, four gRNAs (ACTTGTTCACTGGACCCGCGTGG, TTGTAACAAATGCCGTGGGCTGG, GATGAGGAACGCCTCGGACTGGG, AGAGCATTAAATCTGGCCGTGGG) targeting sequences flanking the critical coding region of Cnpy1 were co-injected with Cas9 mRNA into C57BL/6J zygotes. This strategy produced a precise 6,236 bp deletion, confirmed by PCR and Sanger sequencing.

Genotyping was performed using primers F1 (5’-CTTCAGCCTCTACTCAACAACTTTT-3’) and R1 (5′- CTGTGCGACGTATTCACTCTTTTA-3′), yielding a 490bp product for the knockout allele and a 604bp product for the wild-type allele. A secondary primer set (F2: 5’-TTCCAGATCTATGTGGCACTCCTA-3’; R1 above) was used to distinguish all genotypes: 604 bp (wild-type), 490 bp + 604 bp (heterozygous), and 490 bp (homozygous knockout). Cnpy1 KO mice are viable and fertile.

Heterozygous mice were intercrossed to establish the colony and generate Cnpy1–/– mice for experiments.

#### Additional mouse lines

B6;129P-Tg(Neurog1-cre/ERT2)1Good/J, Strain #:008529 (Koundakjian et al., 2007) and the Rosa26 tdTomato reporters Ai14/B6.Cg-Gt(ROSA)26Sortm14(CAG-tdTomato)Hze/J (Madisen et al., 2010) were purchased from Jackson Laboratory. B6.129S4- Fgfr1tm5.1Sor/J, Strain #:007671 (Hoch and Soriano, 2006) were donated to us by Dr. Pei-San Tsai (University of Colorado Boulder, CO, USA), AP-2εCre/ Tfap2etm1(cre)Will (Feng et al., 2009) (were donated to us by Trevor Williams (University of Colorado Anschutz Medical Campus, Denver, CO, USA), these mice were backcrossed on C57BL/6J for over 10 generations. All experiments using animals were completed in accordance with the guidelines of the Institutional Animal Care and Use Committee (IACUC) at the University at Albany, SUNY.

#### Immunohistochemistry/Immunofluorescence

Mice for immunohistochemistry and immunofluorescence were injected with Sodium Pentobarbital prior to transcardial perfusion with PBS followed by 4% Paraformaldehyde in PBS. Brains were separated from noses and post fixed in 1% Paraformaldehyde overnight at 4°C while noses were post fixed in 4% Paraformaldehyde overnight. Brains were cryoprotected with 30% Sucrose in PBS at 4C overnight and noses of animals older than P21 were incubated in 500mM EDTA in PBS for decalcification. At end of EDTA incubation noses were cryoprotected in sucrose. Samples were embedded in OCT and stored in - 80C prior to cryosectioning serial sections on Superfrost Plus charged slides. Depending on antibody, slides were exposed to antigen retrieval with citric acid buffer prior to blocking and primary antibody incubation at concentration dependent on antibody. Primary antibodies were incubated at 4C overnight. Secondary antibodies were used at 1:1000 concentration and incubated for 1Hr at room temperature. The following primary antibodies were utilized: Chicken Anti-Vglut2 (1:3000, 135416, Santa Cruz), Goat Anti-Kirrel2 (1:500, AF2930, RCD Systems), Mouse Anti-Kirrel3 (1:500, 75-333, NeuroMab), Rabbit Anti-Gαo (1:500, PA5-59337, Invitrogen), Ms Anti-Gαo (1:200, Synaptic Systems 271 111), Mouse Anti-Meis2 63-T (1:500, sc-81986, Santa Cruz), Goat Anti-Ap2ε (1:500, RCD Systems AF5060), Rabbit Anti-Cnpy1 (1:100, NBP2-82720, Novus Bio), Rabit Anti- cCasp3 (1:1000, AB3623, Millipore), Rabbit Anti-Hspa5-488 (1:500, CL488-11587, ProteinTech Group), Rabbit Anti-pS6 Ser 240/244 (1:500, D68F8, Cell Signaling Tech), Rabbit Anti-V2R2 (1:5000, donated by University of Parma). Rabbit Anti-DsRed tomato (1:500, 600-901-379S, Rockland), Goat Anti-DsRed tomato (1:1000, 200-101-379, Rockland), Rabbit Anti-CHOP (1:500, 15204-1-AP, Proteintech).

#### *In-situ* Hybridization

Calr4 double stranded gene block was purchased from Integrated DNA Technologies (IDT) containing the cDNA of interest obtained by Genepaint and were flanked by an upstream T7 promoter and downstream SP6. Hybridizations were performed as described (Lin et al., 2022).

#### Tamoxifen preparation and Use

Tamoxifen (Sigma–Aldrich), CAS # 10540−29−1, was mixed and dissolved in corn oil at a concentration of 20 µg/µl. Pups were injected at 80mg tamoxifen/Kg mouse body weight. Animals were then perfused at the appropriate time point.

#### Male Urine Exposure/pS6 Assay

Mice were weaned, isolated, and habituated in clean static cages for 3 days prior to urine exposure. The mice were placed in a cage saturated with whole-male urine for 90min prior to perfusion. Cells immunoreactive for pS6 above a set threshold were counted to determine activity levels of VSNs in WT mice and Cnpy1 KO mice at age P24.

#### Single Cell mRNA Sequencing

Two animals per genotype were euthanized with CO2 and their VNOs removed and placed immediately in PBS on ice. The non-sensory epithelium was removed from the cartilaginous capsule and digested in Neuronal Isolation Enzyme/Papain, DNAse I, Collagenase A, and L-Cysteine for approximately 15 minutes. Digestion was stopped using 100% FBS and cell suspension filtered with 70μm followed by 40μm cell strainer. Cells were pelleted at 400rcf, 4C, for 5min and resuspended in FBS just prior to 10x protocol.

The 10X Cell Ranger pipeline was used to align FASTQ files and generate Filtered Feature Matrices using the ‘Seurat’ R Package. Principal component analysis (PCA) enables nearest neighbor calculations and clustering of cells based on gene expression, while non-linear dimensional reduction, Uniform Manifold Approximation, and Projection (UMAP) allows for visualization. Cell identity is assigned based on known markers and VSN-specific clusters can be grouped modularly for analysis. The SCTransform Seurat method was used to scale and normalize aligned reads, addressing technical variances in read depth across samples. Clustering resolution was optimized using the Clustree package to clarify cluster boundaries and facilitate targeted exploration of developmental space. Differential Gene Expression (DGE) and representation of UMAPs, Dot plots and heatmaps were performed using the FindMarkers function within the Seurat package utilizing Wilcox ranked sums test. GO Term analysis was performed using the Gost function from the gProfiler package.(Kolberg et al., 2023).

### Quantification, figure preparation, and statistical analysis of microscopy data

Confocal microscopy images were taken on a Zeiss LSM 710 and 980. Epifluorescence images were acquired using a Leica DM4000 B LED fluorescence microscope provided with Leica DFC310 FX camera.

Composite images and figure preparation were prepared using Adobe Photoshop 24.7.0. Prism 10.6.0 was used for all statistical analyses, including calculation of mean values and SEM. Two-tailed, unpaired t-tests were performed for all comparisons, and p-values < 0.05 were considered statistically significant. Sample sizes and p-values are reported as individual data points in each graph and/or in the figure legends. Percentage values were derived following log10 transformation of the data.

#### VNO cell counts and AOB quantifications

For AOB quantifications, established markers were used to determine anterior (Nrp2⁺, Gαi2⁺, Kirrel2+) and posterior (Robo2⁺, Gαo⁺, Kirrel3+) domains of the AOB, and VGLUT2 immunostaining was used to define the glomerular layer. Cell counts obtained by both manual assessment and using a software for automated quantifications that we previously described (Bahreini Jangjoo et al., 2021). For VNO quantifications, manual cell counts were performed on the most medial coronal sections using ImageJ (cell counter) and analyzed as described above. For behavioral experiments, manual cell counts were conducted on confocal or brightfield sections to assess pS6 immunoreactivity in the VNO.

#### Image analysis using Imaris

Confocal image stacks were imported into Imaris (Bitplane) for 3D rendering, mask creation, and fluorescence quantification. Regions of interest were defined by creating surface or volume masks based on a reference marker channel (e.g., ER marker or cell-identity marker), using standardized intensity thresholds and smoothing parameters applied consistently across samples. The fluorescence signal of the target channel was then through the Imaris MeasurementPro module yielding sum of fluorescent intensity and volume for each 3D masked region. To account for variability in mask size, fluorescence values were normalized to mask volume. All parameters (thresholds, filter values, mask creation steps) were kept constant for WT and experimental groups. Data were exported to Excel and then Prism for statistical analysis.

#### Data Sharing

The Cnpy1KO(C57BL/6J-Cnpy1^em1cyagen^ P21 VNO single cell sequencing data is availible available in GEO with accession number <u>GSE310235</u> (https://www.ncbi.nlm.nih.gov/geo/query/acc.cgi?acc=GSE310235). While generated in the same experiment as the Cnpy1 KO data, the Wild-type P21 VNO single cell sequencing data was used for a previous publication(LeFever et al., 2023) and can be found under GEO accession number <u>GSE247872</u>, (https://www.ncbi.nlm.nih.gov/geo/query/acc.cgi?acc=GSE247872).

## Acknowledgments

This work was supported by the Eunice Kennedy Shriver National Institute of Child Health and Human Development (NICHD) under Grants R01HD09733 (P.E.F.) and R01HD114827 (P.E.F.), as well as by the National Institute on Deafness and Other Communication Disorders (NIDCD) under Grant R01DC017149 (P.E.F.). The Zeiss 980 microscope at the University at Albany was funded by the Office of the Director, NIH, under Award Number S10OD028600. We thank Enrico Amato for helping us process the single-cell data and upload it to GEO. We thank Carlsy Ybanez for assisting with Carl4 in-situ hybridization.

